# Differential dynamics specify MeCP2 function at methylated DNA and nucleosomes

**DOI:** 10.1101/2023.06.02.543478

**Authors:** Gabriella N. L. Chua, John W. Watters, Paul Dominic B. Olinares, Joshua A. Luo, Brian T. Chait, Shixin Liu

## Abstract

Methyl-CpG-binding protein 2 (MeCP2) is an essential chromatin-binding protein whose mutations cause Rett syndrome (RTT), a leading cause of monogenic intellectual disabilities in females. Despite its significant biomedical relevance, the mechanism by which MeCP2 navigates the chromatin epigenetic landscape to regulate chromatin structure and gene expression remains unclear. Here, we used correlative single-molecule fluorescence and force microscopy to directly visualize the distribution and dynamics of MeCP2 on a variety of DNA and chromatin substrates. We found that MeCP2 exhibits differential diffusion dynamics when bound to unmethylated and methylated bare DNA. Moreover, we discovered that MeCP2 preferentially binds nucleosomes within the context of chromatinized DNA and stabilizes them from mechanical perturbation. The distinct behaviors of MeCP2 at bare DNA and nucleosomes also specify its ability to recruit TBLR1, a core component of the NCoR1/2 co-repressor complex. We further examined several RTT mutations and found that they disrupt different aspects of the MeCP2-chromatin interaction, rationalizing the heterogeneous nature of the disease. Our work reveals the biophysical basis for MeCP2’s methylation-dependent activities and suggests a nucleosome-centric model for its genomic distribution and gene repressive functions. These insights provide a framework for delineating the multifaceted functions of MeCP2 and aid in our understanding of the molecular mechanisms of RTT.

## Introduction

Methyl-CpG-binding protein 2 (MeCP2) is a highly abundant chromatin-binding protein in mature neurons and is generally thought of as a DNA methylation-dependent transcriptional repressor ^1-3^. Mutations in the X-linked *MECP2* gene cause Rett syndrome (RTT), a severe neurological disorder that occurs 1 in 10,000-15,000 live female births, constituting one of the most frequent causes of monogenic intellectual disabilities in females ^4-6^. Currently there is no known cure for RTT, in part due to the multifaceted and complex functions of MeCP2, which remain poorly understood ^7,8^. As such, elucidating the molecular behavior of MeCP2 and its disease mutants on chromatin is imperative towards establishing therapeutic avenues for targeted intervention.

MeCP2 is a primarily disordered and highly basic protein that exhibits preference for binding methylated cytosines in both CpG and non-CpG contexts but also potently binds unmethylated DNA ^9-11^. In cells, MeCP2 exerts several methylation-dependent functions such as transcriptional repression and transposase protection ^3,12,13^. Additionally, MeCP2 has been shown to interact with and compact nucleosome arrays ^14-16^. However, the pervasive MeCP2 binding sites in the genome and its near-histone-level abundance in neurons has limited our understanding of the preferred chromatin target sites at which MeCP2 performs its function ^8,17^. MeCP2 has also been reported to associate with other effector proteins, most notably the NCoR1/2 co-repressor complex ^18,19^. It is generally presumed that MeCP2’s gene silencing activities are mediated by these effectors, but whether MeCP2 possesses intrinsic properties that enable repression independent of other binding partners remains to be studied. Moreover, how the myriad RTT mutations impair MeCP2’s molecular behavior and function at chromatin remains unclear.

Single-molecule techniques are powerful tools to dissect dynamic and heterogeneous molecular interactions that are difficult to resolve by ensemble methods ^20,21^. In this work, we used correlative single-molecule fluorescence and force microscopy to directly visualize the dynamic interaction of MeCP2 with different types of DNA and chromatin substrates. Our results reveal a remarkably diverse repertoire of binding modes of MeCP2 on chromatin and suggest a nucleosome-centric model for MeCP2’s repressive activities.

## Results

### CpG methylation suppresses MeCP2 diffusion on DNA

We purified recombinant full-length human MeCP2 from *E. coli* and site-specifically labeled the protein with a Cy3 fluorophore (**Figure S1a**). Using a single-molecule instrument that combines dual-trap optical tweezers and confocal fluorescence microscopy ^22^, we first examined the behavior of MeCP2 on methylation-free bacteriophage λ genomic DNA that is 48.5 kilobase pairs (kbp) in length. A single λ DNA molecule was tethered between two optically trapped beads, incubated in a Cy3-MeCP2-containing channel, and then moved to another protein-free channel for imaging (**Figures 1a and S2**). Surprisingly, we observed that MeCP2 often exhibited long-lived and long-range diffusive motions on the DNA (**Figures 1b, c, and S2**). Mean square displacement (MSD) analysis of individual trajectories showed mostly a linear relationship between MSD and the time interval (Δ*t*) (**Figure 1d**), suggesting that MeCP2 undergoes normal Brownian diffusion. The diffusion coefficient (*D*) varied among MeCP2 trajectories (**Figures 1b and e**). Based on the fluorescence intensity of individual trajectories, we found that MeCP2 can bind DNA as multimeric units (8.7 ± 7.9 monomers per trajectory, mean ± SD, *n* = 77) (**Figure S3a**). We plotted *D* against the number of MeCP2 monomers per trajectory and found that larger multimers tend to diffuse slower (**Figure S3b**). Alternatively, local DNA sequences could also contribute to the observed variation in *D* ^23^.

**Figure 1.**
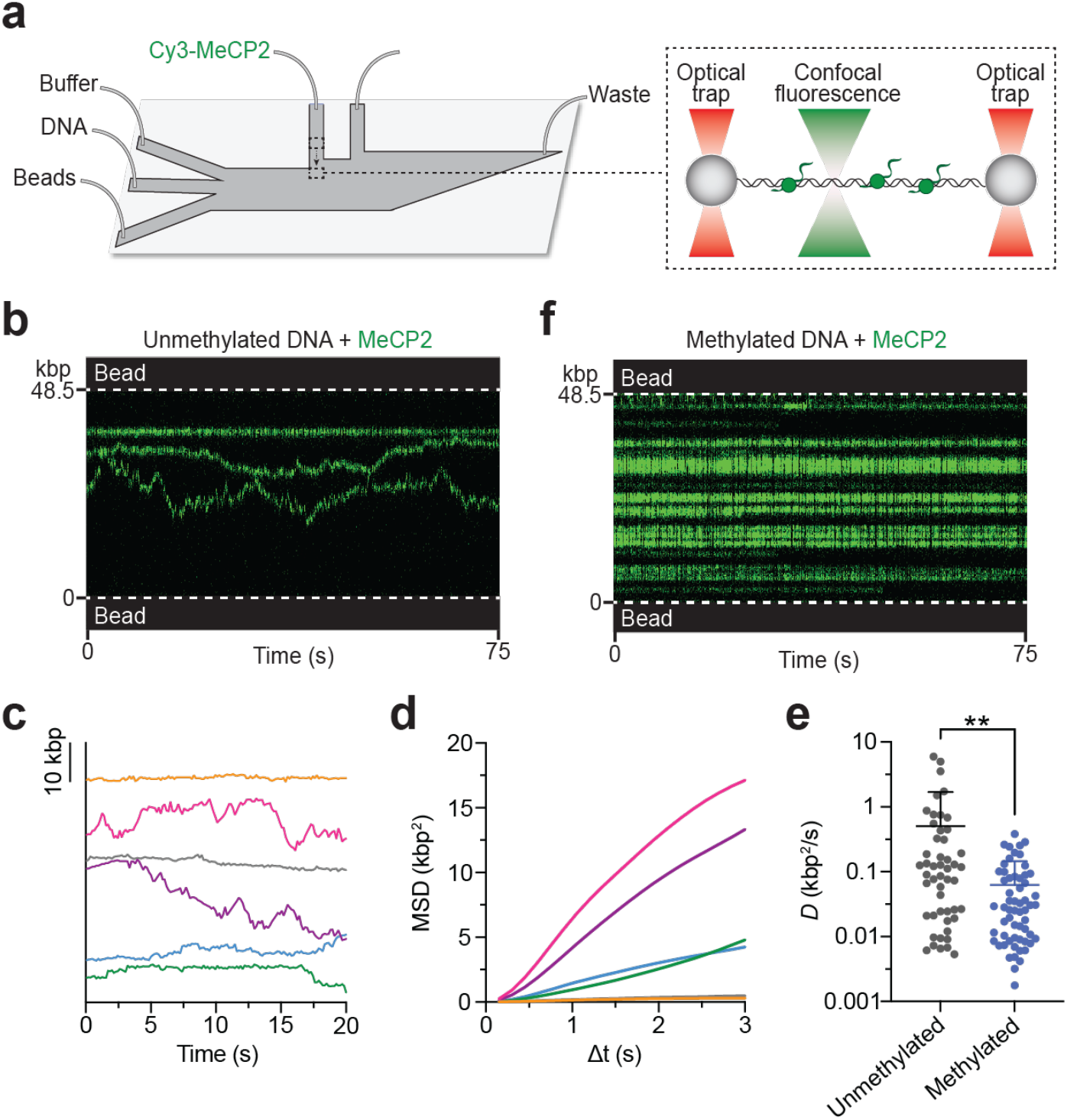
CpG methylation suppresses MeCP2 diffusion on DNA. **a**, Schematic of the experimental setup. A single λ DNA molecule was tethered between a pair of optically trapped beads through biotin-streptavidin linkage. The tether was moved to a channel containing Cy3-MeCP2 to allow protein binding and subsequently to a protein-free channel for imaging. **b**, A representative kymograph of an unmethylated DNA tether bound with Cy3-MeCP2. **c**, Six example MeCP2 trajectories on unmethylated DNA showing their diffusive motions (offset for clarity). **d**, Mean square displacement (MSD) analysis of the trajectories shown in (**c**) (color matched). **e**, Diffusion coefficients (*D*) derived from linear regression of the MSD plots for MeCP2 trajectories on unmethylated or CpG methylated DNA. **f**, A representative kymograph of a methylated DNA tether bound with Cy3-MeCP2.

Next, we used the bacterial M.SssI methyltransferase to methylate the CpG sites within λ DNA (**Figure S1b**) and imaged Cy3-MeCP2 on methylated DNA tethers (**Figure 1f**). Under the same incubation conditions, the average number of MeCP2 trajectories per tether for methylated DNA was significantly higher than that for unmethylated DNA (*k*_on,app_ = 0.37 ± 0.09 s^-1^ and 0.09 ± 0.04 s^-1^, respectively), consistent with bulk results showing that CpG methylation enhances the affinity of MeCP2 to DNA (**Figure S4a**) ^11,16^. Moreover, we observed that methylation substantially suppresses MeCP2 diffusion, which was confirmed by MSD analysis showing a significantly lower average *D* value (**Figures 1e and f**). To exclude the possibility that the suppressed MeCP2 diffusion on methylated DNA was caused by spatial confinement due to enhanced binding, we titrated down the concentration of MeCP2 and observed a similar, static behavior even when the tethers were sparsely bound with MeCP2 (**Figure S4b**). We also plotted the summed MeCP2 signals along the tether length and found that they correlate reasonably well with the distribution of CpG sites on λ DNA (**Figure S4c**). Together, these results reveal that MeCP2 harbors an intrinsic activity to scan on DNA and that such activity is suppressed by CpG methylation.

### RTT mutations differentially perturb MeCP2 behavior on DNA

Our single-molecule platform enabled us to investigate the effects of RTT mutations on MeCP2-DNA interaction. We purified and fluorescently labeled a panel of MeCP2 mutants that display a range of phenotypic severities ^24-26^ (**Figures 2a and S1c**). We first studied T158M, a missense mutation within the methyl binding domain (MBD) of MeCP2 that accounts for ∼12% of all RTT cases ^24,27^. Our data showed MeCP2^T158M^ exhibits significantly reduced binding to methylated DNA compared to the wild-type (WT) protein but no change in its binding to unmethylated DNA (**Figures 2b and S5**), consistent with previous results ^28^. In addition, we also observed reduced diffusion and multimerization on unmethylated DNA for MeCP2^T158M^ compared to WT (**Figures 2c and 2d**).

**Figure 2.**
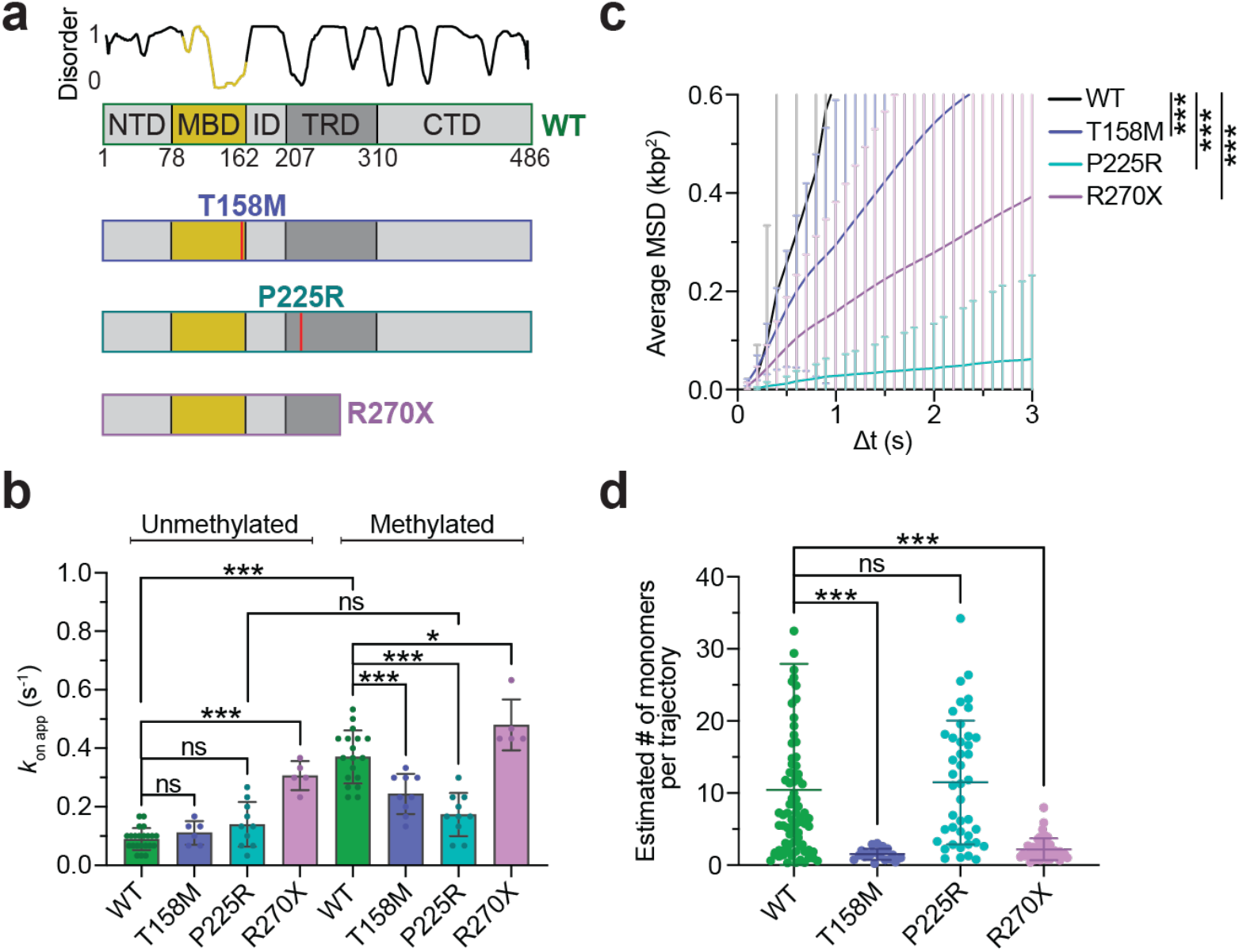
RTT mutations differentially perturb MeCP2 behavior on DNA. **a**, Domain structure of MeCP2 and the corresponding PONDR disorder score. Three RTT mutants (T158M, P225R, and R270X) are shown below the WT protein. **b**, Apparent on rate for WT, T158M, P225R, or R270X MeCP2 binding to unmethylated or methylated DNA. Error bars represent SD. **c**, Average MSD plot for WT (*n* = 78), T158M (*n* = 20), P225R (*n* = 40), or R270X (*n* = 44) Cy3-MeCP2 trajectories on unmethylated DNA. Error bars represent SD. **d**, Estimated number of monomers per trajectory for WT, T158M, P225R, or R270X MeCP2 on unmethylated DNA. Bars represent mean and SD.

Next, we examined the P225R mutation, which resides inside the transcriptional repression domain (TRD) of MeCP2 (**Figures 2a and S1c**). We found that MeCP2^P225R^ exhibits a markedly diminished ability to bind methylated DNA—to an even larger degree than MeCP2^T158M^ (**Figures 2b and S5**). Additionally, P225R drastically slows down MeCP2 diffusion on unmethylated DNA compared to WT (**Figure 2c**). As a result, the behaviors (i.e., binding affinity and diffusivity) of MeCP2^P225R^ on unmethylated versus methylated DNA are indistinguishable, implicating regions outside the MBD in conferring MeCP2 the ability to discriminate between the two forms of DNA.

We then investigated R270X, a truncating mutation lacking the entire C-terminal domain (CTD) and part of the TRD (**Figures 2a and S1c**). Interestingly, MeCP2^R270X^ displayed elevated binding to DNA—especially to the unmethylated form—compared to WT (**Figures 2b and S5**). The truncation also showed significantly reduced diffusion on unmethylated DNA (**Figure 2c**). Moreover, the MeCP2^R270X^ trajectories contained fewer protein monomers on average compared to the WT trajectories (**Figure 2d**), suggesting that the disordered TRD/CTD is at least partially responsible for mediating MeCP2 multimerization on DNA.

### MeCP2 preferentially targets nucleosomes over bare DNA

We next sought to visualize the behavior of MeCP2 on chromatinized DNA. To this end, we reconstituted nucleosomes on unmethylated λ DNA tethers in situ with LD655-labeled human histone octamers using an established protocol ^29^ (**Figure 3a**). Nucleosomes were sparsely loaded (3-10 per tether) so individual loci could be spatially resolved. We then incubated the nucleosomal DNA tether with Cy3-MeCP2 and simultaneously monitored MeCP2 and nucleosome signals via dual-color imaging. Strikingly, we observed frequent colocalization and stable association of MeCP2 with nucleosomes (**Figure 3b**). MSD analysis showed that the nucleosome-associated MeCP2 trajectories were mostly stationary, in stark contrast to the diffusive MeCP2 trajectories located at the intervening bare DNA regions (**Figure 3c**). In the majority of cases, diffusing MeCP2 units were confined between nucleosome-MeCP2 loci and, as a result, their movement was restricted between adjacent nucleosome sites (**Figure 3b**). MeCP2 was also observed to prevalently and stably colocalize with nucleosomes loaded on CpG methylated DNA tethers (**Figure 3d**). As expected, MeCP2 exhibited less diffusion at bare DNA regions within the methylated tethers compared to unmethylated tethers (**Figures 3b, d, and S6**).

**Figure 3.**
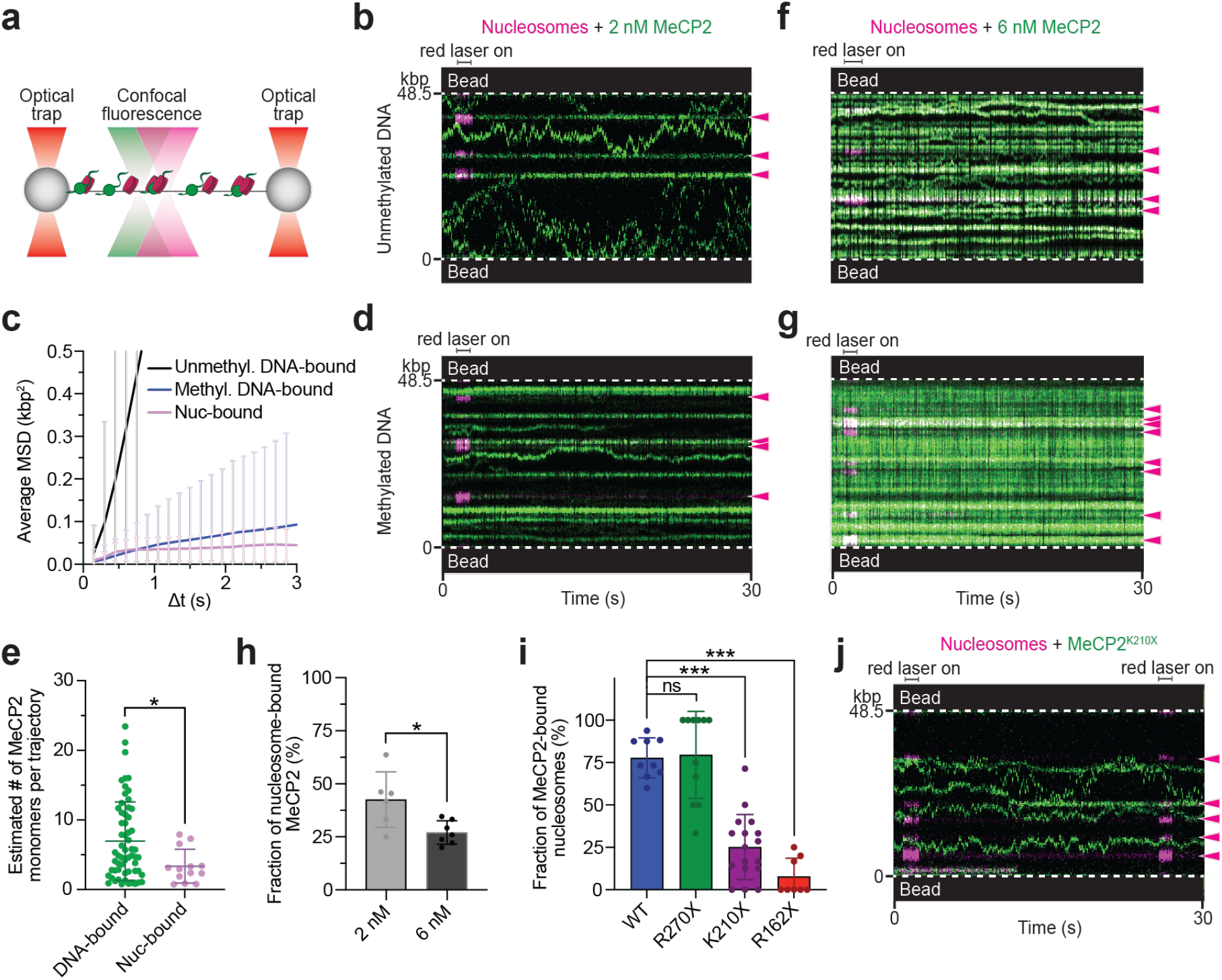
MeCP2 preferentially and stably targets nucleosomes. **a**, Schematic of the experimental setup. Nucleosomes were formed on λ DNA tethers by incubating the tether with LD655-labeled histone octamers and histone chaperone Nap1. LD655-nucleosome and Cy3-MeCP2 signals were simultaneously monitored via two-color confocal microscopy. **b**, A representative kymograph of a nucleosome-containing unmethylated DNA tether incubated with 2 nM Cy3-MeCP2. Red laser was flashed on briefly to locate the nucleosomes within the tether. Arrows denote nucleosome positions. **c**, Average MSD plot for WT Cy3-MeCP2 trajectories on unmethylated DNA (*n* = 78), methylated DNA (*n* = 200), or colocalized with nucleosomes (*n* = 22). Error bars represent SD. **d**, A representative kymograph of a nucleosome-containing methylated DNA tether incubated with 2 nM Cy3-MeCP2. **e**, Estimated number of monomers per trajectory for bare DNA- or nucleosome-bound MeCP2 on unmethylated DNA tethers. Bars represent mean and SD. **f**, A representative kymograph of a nucleosome-containing unmethylated DNA tether incubated with 6 nM Cy3-MeCP2. **g**, A representative kymograph of a nucleosome-containing methylated DNA tether incubated with 6 nM Cy3-MeCP2. **h**, Fraction of Cy3-MeCP2 trajectories on methylated DNA that were colocalized with nucleosomes at 2 nM or 6 nM MeCP2 concentration. The remaining fraction represents MeCP2 bound to bare methylated DNA. Error bars represent SD. **I**, Fraction of nucleosomes on unmethylated DNA tethers that were colocalized with MeCP2 after incubation with 2 nM WT, R270X, K210X, or R162X Cy3-MeCP2. Error bars represent SD. **j**, A representative kymograph of a nucleosome-containing unmethylated DNA tether incubated with 2 nM Cy3-MeCP2^K210X^. Red laser was flashed on periodically to locate the nucleosomes within the tether.

We used native mass spectrometry to examine the interaction between MeCP2 and mononucleosomes and found that they can indeed form stable assemblies (**Figure S7**). We also analyzed the multimeric state of MeCP2 units bound to nucleosomes on λ DNA tethers based on their fluorescence intensities in the kymographs and found that they contained fewer monomers on average than those on bare DNA (**Figure 3e**). Although the number of bare DNA sites greatly outnumbered nucleosome sites on each tether, at 2 nM MeCP2 we observed a comparable number of nucleosome-bound MeCP2 units versus bare-DNA-bound MeCP2 units (**Figures 3b and d**). When we titrated up the MeCP2 concentration used in the experiments, we observed an increasing fraction of bare-DNA-bound MeCP2 units, while the nucleosome sites remained fully occupied (**Figures 3f-h**). These results indicate that nucleosomes serve as preferred target sites for MeCP2 on chromatinized DNA.

To map the MeCP2 domains that are critical for nucleosome binding, we performed single-molecule experiments with a series of MeCP2 truncations: R270X, K210X, and R162X (**Figures S1c and d**). We found that both MeCP2^K210X^ and MeCP2^R162X^ showed significantly diminished nucleosome targeting, whereas MeCP2^R270X^ retained the ability to colocalize with nucleosomes comparable to WT (**Figures 3i, j, and S8**). Therefore, the intervening domain (ID) and part of the TRD (residues 211-270) are critical to MeCP2’s nucleosome-binding activity. Notably, we found that MeCP2^K210X^ and MeCP2^R162X^ still retained the ability to bind and diffuse on bare DNA (**Figures 3j and S8**).

### MeCP2 enhances the mechanical stability of nucleosomes

Next, we explored the functional consequences of MeCP2’s prevalent targeting to nucleosomes. Given their long-lived association, we asked whether MeCP2 binding alters the mechanical properties of the nucleosome. We thus conducted pulling experiments on individual nucleosomal DNA tethers. The resultant force-distance (*F*-*d*) curves contain transitions that signify the unwrapping of individual nucleosomes (**Figure S9**) ^30^. We then repeated the pulling experiments in the presence of MeCP2 and simultaneously monitored the fluorescence signals from Cy3-MeCP2 and LD655-nucleosomes. MeCP2 was observed to mostly remain associated with the nucleosomes throughout pulling (**Figure 4a**). We then analyzed the *F*-*d* curves and found that MeCP2 significantly increased the average force required to unwrap nucleosomes (**Figures 4b and c**), providing evidence for a direct stabilization effect of MeCP2 on the nucleosome. Interestingly, this effect was diminished when the WT MeCP2 was replaced with MeCP2^R270X^ (**Figure 4c**), even though the truncated and WT proteins displayed a similar level of nucleosome binding (**Figure 3i**).

**Figure 4.**
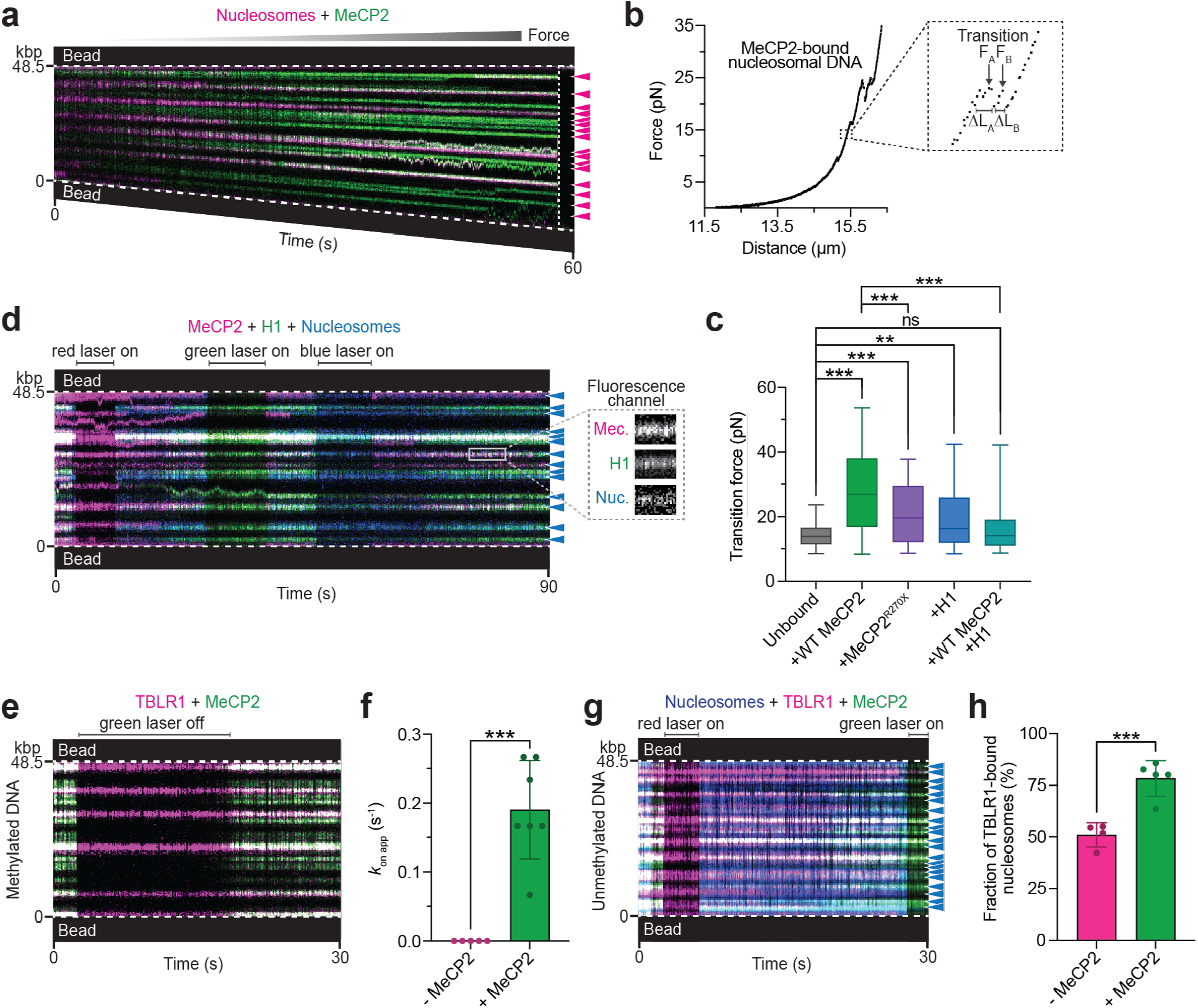
MeCP2 exerts stabilization and recruitment functions at nucleosomes. **a**, A representative kymograph of a nucleosome-containing unmethylated DNA tether bound with Cy3-MeCP2 and pulled to high forces by gradually increasing the inter-bead distance. Vertical dotted line denotes the time when the tether ruptured. Arrows denote nucleosome positions. **b**, A representative force-distance curve of a MeCP2-bound nucleosome-containing DNA tether showing force-induced nucleosome unwrapping transitions. Inset shows a zoom-in view of two example transitions for which the distance changes (Δ*L*) and the transition forces are recorded. **c**, Distribution of transition forces recorded from force-distance curves of nucleosomal DNA tethers with no MeCP2 or H1 bound (*n* = 84), or bound with WT MeCP2 (*n* = 107), MeCP2^R270X^ (*n* = 68), H1 (*n* = 106), or with both WT MeCP2 and H1 (*n* = 81). Box boundaries represent 25^th^ to 75^th^ percentiles, middle bar represents median, and whiskers represent minimum and maximum values. **d**, A representative kymograph of an unmethylated DNA tether containing AF488-labeled nucleosomes and incubated with Cy5-labeled MeCP2 and Cy3-labeled H1. Lasers were alternated on and off to confirm signal from each fluorescence channel. Arrows denote nucleosome positions. Inset shows a zoom-in view of individual fluorescence channels at a nucleosome site where both MeCP2 and H1 colocalized. **e**, A representative kymograph of a methylated DNA tether incubated with Cy3-labeled MeCP2 and LD655-labeled TBLR1. **f**, Apparent on rate for TBLR1 binding to methylated DNA in the absence or presence of MeCP2. Error bars represent SD. **g**, A representative kymograph of an AF488-nucleosome-containing unmethylated DNA tether incubated with Cy3-MeCP2 and LD655-TBLR1. Arrows denote nucleosome positions. **h**, Fraction of nucleosomes on unmethylated DNA tethers that were colocalized with TBLR1 in the absence or presence of MeCP2. Error bars represent SD.

### MeCP2 and H1 colocalize with nucleosomes

Linker histone H1 is a major component of eukaryotic chromatin that also binds and organizes nucleosomes ^31^. The interplay between MeCP2 and H1 in chromatin regulation is under debate, although some literature has suggested that MeCP2 and H1 antagonize each other for nucleosome interaction ^1,3,32,33^. To directly visualize their behaviors on chromatin, we performed three-color single-molecule fluorescence experiments with Cy3-H1.4, Cy5-MeCP2, and AF488-labeled nucleosomes loaded on unmethylated DNA tethers. Contrary to an antagonistic binding model, we observed frequent colocalization of H1 and MeCP2 at nucleosome sites—MeCP2 signal was detected at 63% of H1-bound nucleosomes (**Figures 4d and S10a**). We then investigated how the co-binding of MeCP2 and H1 impinges on nucleosome stability. Significantly, pulling on nucleosomal DNA tethers incubated with both MeCP2 and H1.4 yielded an average transition force lower than the value for nucleosomes incubated with only WT MeCP2 and similar to the value for unbound nucleosomes (**Figures 4c and S10b**). These results reveal that the nucleosome can simultaneously accommodate both H1 and MeCP2, but H1 alleviates the nucleosome stabilization effect of MeCP2, which indicates that the binding pose of MeCP2 in the ternary complex is distinct from that of MeCP2 bound to the nucleosome alone.

### MeCP2 specifies TBLR1 recruitment to methylated DNA and nucleosomes

Finally, we investigated the recruitment function of MeCP2 in light of its differential behaviors on DNA and nucleosomes. It has been reported that MeCP2 associates with the NCoR1/2 co-repressor complex through its transducing-beta like 1-related (TBLR1) core components ^18,19,34^ and recruits this complex to methylated heterochromatin ^25^. To resolve the interaction between MeCP2 and TBLR1 at specific chromatin substrates, we performed single-molecule experiments to track the behaviors of LD655-labeled CTD domain of TBLR1 (TBLR1^CTD^) and Cy3-MeCP2 on bare DNA and nucleosomal DNA tethers. We found that the recruitment of TBLR1 to bare methylated DNA is strictly dependent on MeCP2—no TBLR1 signal was detected on the DNA in the absence of MeCP2, while stable TBLR1 trajectories were observed when MeCP2 was present and the two proteins always colocalized (**Figures 4e, f, and S10c**). TBLR1 can also be recruited by MeCP2 to bare unmethylated DNA, albeit at a lower frequency (**Figures S10d and e**). Notably, TBLR1 was occasionally observed to be co-diffusing with MeCP2 along unmethylated DNA (**Figure S10d**). With nucleosomal DNA, we found that TBLR1 readily binds nucleosomes alone (**Figure S10f**), but MeCP2 significantly increased the frequency of TBLR1 recruitment to nucleosome loci (**Figures 4g and h**). These findings suggest that MeCP2 directs TBLR1 recruitment to chromatin through its distinct DNA- and nucleosome-binding modes.

## Discussion

More than two decades after the discovery of mutations in MeCP2 as the genetic drivers of RTT, the biophysical and biochemical properties of this protein remain to be fully characterized. In particular, its dynamics and distribution on individual chromatin substrates have not been studied, which likely underlie the multiplexed functions of this abundant chromatin-binding protein. In this study, we used single-molecule visualization and manipulation to dissect the behavior of full-length MeCP2 and its mutants on DNA and chromatin. First, we discovered that MeCP2 can quickly scan DNA via one-dimensional diffusion, a property shared by other DNA-binding proteins and understood to facilitate target search ^35,36^. MeCP2 diffusion is greatly suppressed by CpG methylation, which poises the protein to mediate methylation-dependent activities such as NCoR1/2 recruitment (**Figure 5**). The different kinetic nature of MeCP2 on unmethylated versus methylated DNA also rationalizes how MeCP2 protects methyl CpG sites from transposase ^13^, MNase ^15^, and DNaseI ^37^ digestion, while retaining a comparable binding affinity to unmethylated DNA ^11^. Notably, we found that some RTT mutations, such as P225R, cause much reduced diffusion on unmethylated DNA. It is conceivable that the pathological mechanism for these patients may be related to ectopic MeCP2 activities that are normally restricted among methylated DNA sites.

**Figure 5.**
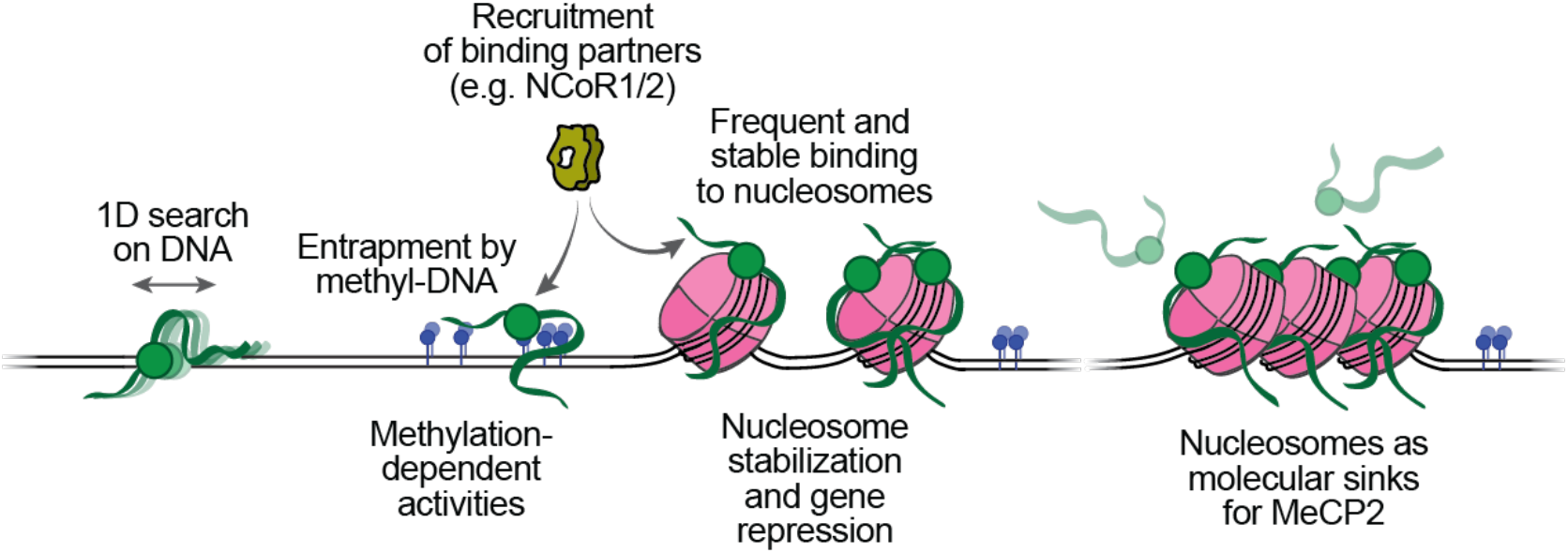
Working model for MeCP2 functioning at chromatin. MeCP2 rapidly scans unmethylated bare DNA regions and becomes trapped upon encountering methyl-DNA sites, where it performs methylation-dependent activities such as recruiting transcriptional co-repressors. In contrast, MeCP2 stably engages with nucleosomes and protects them from mechanical perturbation. This interaction also facilitates the recruitment of binding partners to nucleosome sites. Finally, nucleosomes capture the majority of MeCP2 molecules in the nucleus, leaving only a fraction of free proteins to bind bare DNA. This provides a plausible explanation for why even a modest change in the MeCP2 level can drastically alter its regulatory function. Therefore, MeCP2 plays both direct and indirect roles in chromatin organization and gene regulation dependent on its differential dynamics at various regions of the genome.

Our study also provides insights into the behavior of MeCP2 in the chromatin context. Echoing previous biochemical results showing that MeCP2 can directly bind nucleosomes ^14-16^, our single-molecule results further reveal that the MeCP2-nucleosome interaction is prevalent and stable, contrasting the protein’s dynamic behavior on bare DNA. We propose that this stable association mediates hitherto underappreciated nucleosome-directed activities of MeCP2 (**Figure 5**). In support of this notion, we demonstrate that MeCP2 binding alone stabilizes nucleosomes against mechanical unwrapping, which could suppress the activities of chromatin remodelers and the transcription machinery. Intriguingly, we found the co-binding of H1 attenuates this stabilizing effect. Thus, the presence of H1, another abundant chromatin-binding protein, antagonizes the activity of MeCP2 through mechanical regulation rather than competitive binding to the nucleosome. We also show that MeCP2 enhances the binding of TBLR1 to nucleosomes, suggesting another mechanism to functionalize the MeCP2-nucleosome interaction. These findings are compatible with the previously proposed “bridge hypothesis,” which postulates that MeCP2 recruits the NCoR1/2 co-repressor complex to methylated heterochromatin to execute its role as a global repressor ^17^. Our single-molecule results further sharpen this model by demonstrating this recruitment function occurs primarily at methylated DNA and nucleosome sites. Considering the near stoichiometric amount of MeCP2 to histones in neuronal nuclei, MeCP2 likely recruits other effectors to nucleosome sites. The current work establishes an experimental platform to screen the candidates and directly examine their interplay with MeCP2.

We show that MeCP2 frequently targets nucleosomes even in the presence of many more available DNA binding sites. Unlike the sparsely loaded DNA used in our experiments, DNA inside the nucleus is predominantly wrapped into nucleosomes. Therefore, it is likely that nucleosomes capture the majority of MeCP2 in vivo, leaving only a small fraction of proteins to bind bare DNA (**Figure 5**). This model provides a plausible explanation for why even a modest change in the MeCP2 level can drastically alter its regulatory function. Indeed, it is known that neurons are highly sensitive to the MeCP2 dosage, and both mild under- and over-expression can lead to disease ^24,25,38,39^. Considering that the relative levels of nucleosome-versus DNA-bound MeCP2 may be important for maintaining their respective functions, we found that the region between K210 and R270 is crucial for MeCP2’s nucleosome-binding activity but not for its ability to bind DNA, indicating RTT truncating mutations in this area (S204X, G232fs, R255X, etc.) may shift the proper balance of MeCP2 distribution, resulting in abnormal function. It will be important to investigate if histone modifications, variants, and other chromatin-binding proteins also modulate the distribution of MeCP2 among different regions of the chromatin.

In sum, our study reveals that MeCP2 differentially interacts with DNA and nucleosomes, allowing it to serve distinct biophysical and recruiting roles. RTT mutations alter these interactions in different ways and to different degrees. These insights will help develop targeted intervention strategies to restore the normal functioning of MeCP2 at chromatin.

## Methods

### Protein purification and fluorescent labeling

#### MeCP2

Human MeCP2 in the pTXB1 plasmid (Addgene #48091) was propagated in *E. coli* 5-alpha cells (New England BioLabs). Following mutagenesis for fluorescent labeling and/or creating RTT mutations using the Q5 mutagenesis kit (New England BioLabs), plasmids were transformed into *E. coli* BL21(DE3) cells (Thermo Fisher) for overexpression. Expression and purification of MeCP2 was achieved by starting with chitin-intein MeCP2 fusion proteins. The protocol was adapted from a previously published protocol ^40^ and the manufacturer’s instructions for the IMPACT system (New England BioLabs). 4 L of cells in the presence of 100 μg/mL carbenicillin were grown to an OD_600_ of 0.5 and induced with 0.5 mM IPTG overnight at 16°C. Lysates were prepared by resuspending cell pellets in column buffer (20 mM Tris hydrochloride pH 8.0, 500 mM sodium chloride, 0.1% Triton X-100, and 0.1 mM PMSF [GoldBio]) followed by sonication and centrifugation at 14,000 rpm for 30 min. Lysates were applied to 10-mL bed volume of chitin resin (New England BioLabs) that was pre-equilibrated with column buffer for 1.5 hours at 4°C on a tube rotator. The resin was washed with 20× resin bed volumes of column buffer and then flushed with 3× resin bed volumes of column buffer supplemented with 50 mM dithiothreitol before being capped and left overnight at room temperature for intein cleavage. Fractions were eluted with column buffer and analyzed by SDS-PAGE, and peak fractions were pooled, concentrated, and added to a Superdex 200 Increase 10/300 GL column equilibrated with column buffer attached to an AKTA pure system (Cytiva) for gel filtration. Peak fractions were analyzed by SDS-PAGE and aliquoted for fluorescent labeling or flash frozen and stored in -80°C. To obtain WT MeCP2 labeled with a single Cy3 or Cy5 fluorophore, 2 out of 3 cysteine residues were mutated to serine (C339S, C413S), leaving a single cysteine residue (C429) located towards the end of the disordered CTD. None of the labeling positions used have been implicated in RTT. C429 WT MeCP2 was expressed and purified as described and subsequently incubated with 3× molar excess of tris carboxy ethyl phosphene (TCEP) at 4°C for 30 min. Cy3- or Cy5-maleimide dye (Cytiva) was added to achieve a 10:1 molar ratio of dye to MeCP2 and incubated at 4°C overnight in the dark. To remove free dye, labeled protein was dialyzed in 3× 1-L column buffer and subsequently analyzed by SDS-PAGE, concentrated, aliquoted, flash frozen, and stored in -80°C. The final labeling efficiency was estimated to be ∼80%. A similar protocol was performed to fluorescently label MeCP2 mutants, several containing different labeling positions and associated labeling efficiencies ranging from 80-100%: Cy3-C429 T158M MeCP2, Cy3-C429 P225R MeCP2, Cy3-S242C 270X MeCP2, Cy3-S194C K210X MeCP2, and Cy3-S13C R162X MeCP2.

#### Histone Octamers

Recombinant human core histones were purified and labeled with an LD655 or AF488 fluorophore as previously described ^41^. Briefly, core histones and their labeling mutants were individually expressed in *E. coli* BL21 (DE3) cells, extracted from inclusion bodies and purified under denaturing conditions using Q and SP ion exchange columns (GE Healthcare). H4^L50C^ was labeled with LD655-maleimide (Lumidyne Technologies) under denaturing conditions. Octamers were reconstituted by adding equal ratios of each core histone (H4^LD655-L50C^, H3.2, H2A, and H2B) and purified by gel filtration as described previously. The same protocol was performed to obtain histone octamers containing AF488-H2A^K12C^.

#### TBLR1

Recombinant human TBLR1^CTD^ (residues 134-514) was inserted into a pCAG-TEV-3C plasmid, and the GFP-fusion protein was expressed in 400 mL of suspension HEK293 cells. Cell pellet was lysed in 20-mL lysis buffer (50 mM Tris hydrochloride pH 8.0, 300 mM sodium chloride, 3 mM 2-mercaptoethanol, 0.2% NP40, 1 mg/mL aprotinin, 1 mg/mL leupeptin, 1 mg/mL pepstatin A, 100 mM PMSF, 2 mM ATP, and 2 mM magnesium chloride) with the addition of 1 μL Benzonase (Millipore Sigma) by vortexing. The solution was nutated on a rotating nutator at 4°C for 20 min and centrifuged at 20,000 rpm for 30 min at 4°C. The lysate was collected, added to 1 mL of GFP nanobody coated agarose bead slurry that was pre-equilibrated with wash buffer (50 mM Tris hydrochloride pH 8.0, 300 mM sodium chloride, and 3 mM 2-mercaptoethanol), and nutated on a rotating nutator at 4°C for 1.5 hours. The beads were pelleted by centrifugation at 1000× g for 2 min and the supernatant was removed. The beads were washed 3× with 1 mL wash buffer to remove detergent and protease inhibitors. The beads were resuspended in 250 μL wash buffer and 250 μL of 3C protease was added. The bead solution was nutated at 4°C in the rotating nutator overnight. Beads were then pelleted by centrifugation at 1000× g for 2 min at 4°C and the supernatant was collected. This step was repeated 3× after the addition of wash buffer to collect 5 mL of supernatant total. The eluted protein was then concentrated and added to a Superdex 200 Increase 10/300 GL column equilibrated with wash buffer attached to an AKTA pure system (Cytiva) for gel filtration. Peak fractions were analyzed by SDS-PAGE and aliquoted for fluorescent labeling.

To non-specifically attach a fluorophore to the N terminus of TBLR1^CTD^, the purified protein was dialyzed in 3× 1 L of labeling buffer (45 mM HEPES pH 7.0, 200 mM sodium chloride, 1 mM dithiothreitol, and 0.25 mM EDTA) and LD655-NHS dye (Lumidyne Technologies) was added to achieve a 5:1 molar ratio of dye to TBLR1^CTD^. The mixture was incubated at room temperature for 1 hour in the dark, and the reaction was quenched by adding 30 mM Tris hydrochloride pH 7.0 for 5 min at room temperature. To remove free dye, labeled protein was dialyzed in 3× 1 L of storage buffer (45 mM HEPES pH 7.6, 200 mM sodium chloride, and 1 mM dithiothreitol) and subsequently analyzed by SDS-PAGE, concentrated, aliquoted, flash frozen, and stored in -80°C. The final labeling efficiency was estimated to be ∼85%.

Recombinant *S. cerevisiae* Nap1 was expressed and purified as previously described ^29^.

Recombinant linker histone H1.4^A4C^ was purified and labeled with a Cy3 fluorophore as previously described ^42^.

### DNA substrate preparation

#### Biotinylated DNA

To generate terminally biotinylated λ genomic double-stranded DNA, the 12-base overhang on each end of Dam and Dcm methylation-free *E. coli* bacteriophage DNA (48,502 bp; Thermo Fisher) was filled in with a mixture of natural and biotinylated nucleotides by the exonuclease-deficient DNA polymerase I Klenow fragment (New England BioLabs). The reaction was performed by incubating 17 μg λ DNA, 32 μM each of dGTP/biotin-14-dATP/biotin-11-dUTP/biotin-14-dCTP (Thermo Fisher), and 5 U Klenow in 1× NEBuffer 2 (New England BioLabs) (120 μL total volume) at room temperature for 15 min. The reaction was stopped by adding 10 mM EDTA and heat inactivated at 75°C for 20 min. Biotinylated DNA was then ethanol precipitated for at least 1 hour at -20°C in 3× volume ice-cold ethanol and 300 mM sodium acetate pH 5.2. Precipitated DNA was recovered by centrifugation at 12,000 rpm for 30 min at 4°C. After removing the supernatant, the pellet was washed twice with 1 mL of 70% ethanol, each round followed by centrifugation at 12,000 rpm for 1 min at 4°C and removal of the supernatant. The resulting pellet was air-dried, resuspended in TE buffer (10 mM Tris-HCl pH 8.0, 1 mM EDTA), and stored at 4°C.

#### CpG methylated DNA

To generate CpG methylated DNA, 500 ng biotinylated λ DNA was incubated at 37°C with 1.6 M S-adenosylmethionine (New England BioLabs) and 20 U CpG methyltransferase M.SssI (New England BioLabs) (20 μL total volume) overnight. The reaction was stopped by heat inactivation at 65°C for 20 min. Methylation efficiency was assessed by incubating methylated DNA with the CpG methylation-sensitive restriction enzyme, BstUI (New England BioLabs), which is unable to perform digestion in the presence of methylation at its cut site (157 predicted sites on λ DNA).

### Single-molecule experiments

#### Experimental setup

Single-molecule experiments were performed at room temperature on a LUMICKS C-Trap instrument, which combines three-color confocal fluorescence microscopy with dual-trap optical tweezers ^43^. Rapid optical trap movement was enabled by a computer-controlled stage within a five-channel flow cell (**Figure 1a**). Channels 1-3 were separated by laminar flow, which were used to form DNA tethers between two 3.13-μm streptavidin-coated polystyrene beads (Spherotech) held in traps with a stiffness of ∼0.6 pN/nm. Under a constant flow, a single bead was caught in each trap in channel 1. The traps were then moved to channel 2, and biotinylated DNA was caught between both traps, as detected by a change in the force-distance curve. Flow was stopped and the traps were moved into channel 3 containing only buffer where the presence of a single DNA tether was confirmed by the force-distance curve. Channels 4 and 5 were loaded with proteins as described for each assay. Flow was turned off during data acquisition and visualization of protein behavior.

#### Fluorescence detection

Cy3, LD655, and AF488 fluorophores were excited by three laser lines at 532, 638, and 488 nm respectively. Kymographs were generated by confocal line scanning through the center of the two beads at 100 ms/line. Individual lasers were occasionally turned off to confirm the presence of other fluorophore-labeled proteins. To investigate the behavior of Cy3-MeCP2 on DNA, optical traps tethering a λ DNA molecule under 1 pN of constant tension were moved into channel 4 of the microfluidic flow cell containing 2 nM of Cy3-MeCP2 (unless specified otherwise) in an imaging buffer containing 20 mM Tris hydrochloride pH 8.0 and 100 mM sodium chloride. Following 30-s incubation, the tether was moved to channel 3 containing buffer only for removal of nonspecific binding events and imaging.

To generate nucleosome-containing DNA tethers, optical traps tethering a λ DNA molecule under 1 pN of constant tension were moved into channel 4 of the microfluidic flow cell containing 2 nM LD655-histone octamers and 2 nM Nap1 in 1× HR buffer (30 mM Tris acetate pH 7.5, 20 mM magnesium acetate, 50 mM potassium chloride, and 0.1 mg/mL BSA). Following a 3-s incubation (both octamer concentration and incubation time were optimized to form 3-10 nucleosomes on each DNA tether), tethers were moved to channel 3 containing 0.25 mg/mL sheared salmon sperm DNA in 1× HR buffer for removal of nonspecific octamer binding. Formation of properly wrapped nucleosomes was confirmed by pulling the tether to generate force-distance curves showing force-induced transitions of expected distance change occurring at expected force regime ^44^.

To investigate the behavior of Cy3-MeCP2 on this substrate, a nucleosome-containing DNA tether was moved to channel 5 containing 2 nM of Cy3-MeCP2 (unless specified otherwise) in imaging buffer. Following a 30-s incubation, the tether was moved to channel 3 for imaging.

To investigate the interplay between MeCP2 and H1, AF488-nucleosome-containing DNA tethers were moved to channel 5 containing 2 nM of LD655-MeCP2 and 10 nM of Cy3-H1 in imaging buffer. Following a 30-s incubation, tethers were moved to channel 3 for imaging.

The same protocol was used to investigate the interplay between MeCP2 and TBLR1, except that 2 nM of Cy3-MeCP2 and 20 nM of LD655-TBLR1^CTD^ were used.

#### Force manipulation

Nucleosomal DNA tethers (unbound or bound with MeCP2/H1 proteins) were first relaxed by lowering the distance between traps in channel 3 until ∼0.25 pN of force was reached. The force was zeroed, and the tether was subjected to pulling by moving one trap relative to the other at a constant velocity of 0.1 μm/s until DNA entered the over-stretching region (60-65 pN) or the tether broke.

#### Data analysis

Kymographs were processed and analyzed using a custom script (https://harbor.lumicks.com/single-script/c5b103a4-0804-4b06-95d3-20a08d65768f) which incorporates tools from the lumicks.pylake Python library and other Python modules (Numpy, Matplotlib, Pandas) to generate tracked lines using the kymotracker greedy algorithm.

To determine the mean squared displacement (MSD), the tracked lines were then smoothed using a 3^rd^ order Savitzky-Golay filter with a window length of 11 tracked frames, and the MSD was calculated from each smoothed line. The diffusion coefficient (*D*) was calculated by fitting the MSD trajectory to the equation for 1D diffusion where *MSD* = 2*Dt*^∝^ [alpha is the exponential term used to characterize normal diffusion (*α* = 1), sub-diffusion (*α* < 1), or super-diffusion (*α* > 1)]. The fit was discarded if the *R*^2^ value of the fit was less than 0.8.

The estimated number of monomers per trajectory was determined by dividing the photon count for each trajectory averaged over a 30-s time window by the photon count for a single Cy3-MeCP2 or LD655-nucleosome excited in our instrument.

The apparent on rate of MeCP2 or TBLR1 to DNA was calculated as the number of fluorescence trajectories per tether divided by the incubation time in the protein channel. Only stably bound proteins were considered, defined as those that survived longer than 30 s in the protein-free buffer channel.

Force-distance curves for nucleosome unwrapping experiments were analyzed by extracting the distance change (Δ*L*) and the transition force of abrupt rips associated with individual nucleosome unwrapping events. Only rips above 8 pN were analyzed, which correspond to unwrapping of the inner DNA turn of the nucleosome ^44^.

### Native mass spectrometry (nMS) analysis

2 μM of the reconstituted nucleosome was mixed with MeCP2 at varying molar ratios and then buffer-exchanged into nMS solution (150 mM ammonium acetate, pH 7.5, 0.01% Tween-20) using Zeba desalting microspin columns with a 40-kDa molecular weight cut-off (Thermo Scientific). Each nMS sample was loaded into a gold-coated quartz capillary tip that was prepared in-house and was electrosprayed into an Exactive Plus EMR instrument (Thermo Fisher Scientific) using a modified static nanospray source ^45^. The MS parameters used included: spray voltage, 1.22 kV; capillary temperature, 150 °C; S-lens RF level, 200; resolving power, 8,750 at m/z of 200; AGC target, 1 × 10^6^; number of microscans, 5; maximum injection time, 200 ms; in-source dissociation (ISD), 0 – 10 V; injection flatapole, 8 V; interflatapole, 4 V; bent flatapole, 4 V; high energy collision dissociation (HCD), 150 – 180 V; ultrahigh vacuum pressure, 5 × 10^−10^ mbar; total number of scans, 100. Mass calibration in positive EMR mode was performed using cesium iodide. Raw nMS spectra were visualized using Thermo Xcalibur Qual Browser (version 4.2.47). Data processing and spectra deconvolution were performed using UniDec version 4.2.0 ^46,47^.

Native MS analysis of the four individual histone proteins and MeCP2 confirmed their primary sequence and revealed that these proteins had undergone canonical N-terminal processing (removal of N-terminal methionine). In addition, unbound bacterial DnaK was observed in the MeCP2 sample. Overall, the following expected masses based on the sequence after N-terminal processing were used for the component proteins— H2A: 13,974.3 Da; H2B: 13,758.9 Da; H3.2: 15,256.8 Da; H4.A: 11,236.1 Da; MeCP2: 52,309.4 Da. Based on its sequence, the mass of the 207-bp dsDNA used was 127,801.8 Da. For the reconstituted nucleosome sample, we obtained one predominant peak series corresponding to the fully assembled nucleosome (histone octamer + dsDNA) with a measured mass of 236,277 Da (mass accuracy of 0.01%). Addition of two-fold or five-fold molar excess of MeCP2 to the nucleosome sample yielded additional peaks that corresponded to the binding of one or two MeCP2 (**Figure S7**). Mononucleosomes used for nMS were assembled by salt gradient dialysis using unlabeled WT human histone octamers as previously described ^48^.

### Electrophoretic mobility shift assay (EMSA)

5 nM of 147-bp unmethylated or CpG methylated DNA was incubated with the indicated concentration of WT MeCP2 at room temperature for 5 min with imaging buffer (20 mM Tris hydrochloride pH 8.0 and 100 mM sodium chloride) in a total volume of 10 μL. 1.5 μL of 2 M sucrose was added and 10 μL of each sample was run on a 5% native PAGE gel at 110 V for 60 min on ice. The DNA was stained with SYBR Safe DNA Gel Stain (Thermo Fisher) and visualized using a gel imager (Axygen).

### Statistical analysis

Errors reported in this study represent the standard deviation. *P* values were determined from two-tailed two-sample t tests (*p < 0.05; **p < 0.01; ***p < 0.001) for Figure 1e, Figure 3e, h, Figure 4f, and h. *P* values were determined from a one-way ANOVA with Tukey’s test for multiple comparisons (*p < 0.05; **p < 0.01; ***p < 0.001) for all other statistical analyses.

## Acknowledgments

We thank V. Risca, H. Funabiki, and Y. Arimura for their advice on the project. We thank R. Gong, D. Phua, and G. Alushin for help with TBLR1 purification. G.N.L.C. acknowledges support from the National Institute of Mental Health of the National Institutes of Health (NIH) under award number F31MH132306. J.W.W. is supported by an NRSA Training Grant from the NIH (T32GM066699). B.T.C. acknowledges support from the NIH under award numbers P41GM109824 and P41GM103314. S.L. is supported by an Innovation Award from the International Rett Syndrome Foundation, the Robertson Foundation, the Pershing Square Sohn Cancer Research Alliance, and an NIH Director’s New Innovator Award (DP2HG010510).

## Author contributions

G.N.L.C. conceived the project, prepared the reagents, designed and performed the single-molecule and bulk experiments. J.W.W. wrote the scripts for and assisted with single-molecule data analysis. P.D.O. and B.T.C. performed the native mass spectrometry experiments. J.A.L. assisted with the preparation of several RTT mutant proteins and single-molecule experiments. S.L. oversaw the project. G.N.L.C. and S.L. wrote the manuscript.

