## Supplemental Figures for "Differential dynamics specify MeCP2 function at methylated DNA and nucleosomes"

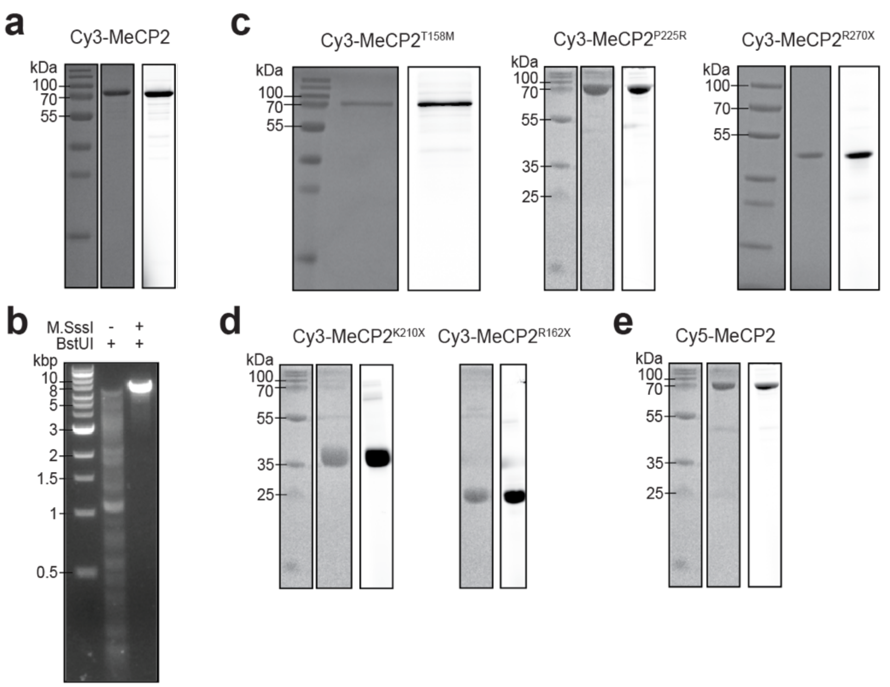


**Figure S1. Evaluation of reagents used in this work.**

**a**, The purity of Cy3-labeled WT MeCP2 was analyzed by SDS-PAGE via Coomassie Blue staining (middle) and fluorescence scanning (right). Protein ladder is shown on the left. **b**, CpG methylation by M.SssI bacterial methyltransferase is assessed via BstUI digestion. Agarose gel (1%) shows blocked BstUI digestion of a 9-kbp linear DNA after M.SssI treatment. **c**-**e**, The purity of Cy3-MeCP2$^{T158M}$, Cy3-MeCP2$^{P225R}$, Cy3-MeCP2$^{R270X}$ (**c**), Cy3-MeCP2$^{K210X}$, Cy3-MeCP2$^{R162X}$ (**d**), and Cy5-labeled WT MeCP2 (**e**) was analyzed by SDS-PAGE via Coomassie Blue staining and fluorescence scanning.

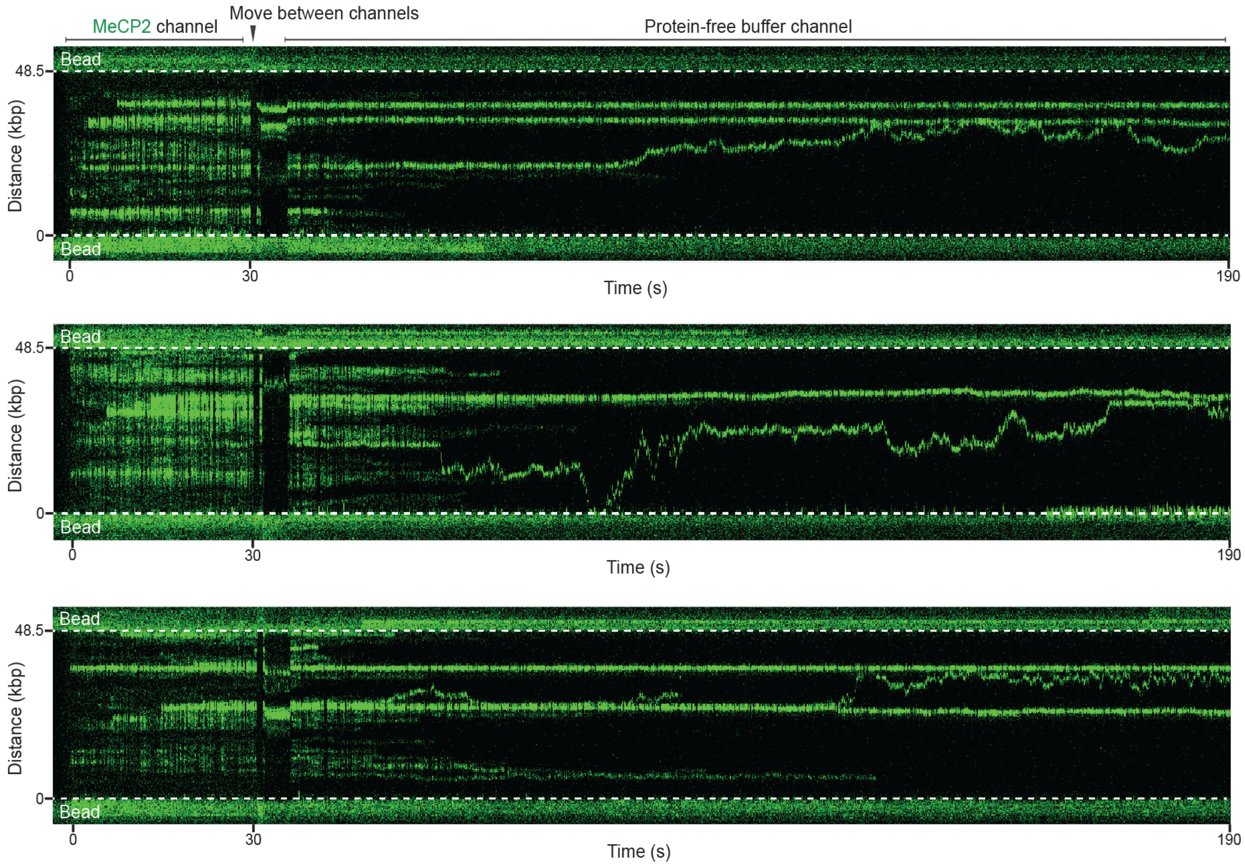


**Figure S2. Additional kymographs of unmethylated DNA tethers bound with Cy3-MeCP2.**

Each λ DNA tether was incubated in a channel containing 2 nM Cy3-MeCP2 for 30 s and then moved to a protein-free buffer channel for imaging.

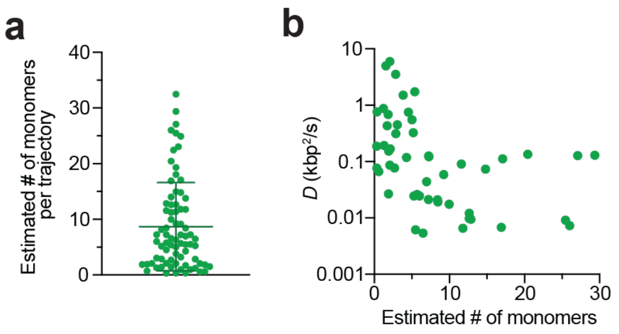


**Figure S3. Analysis of MeCP2 multimerization on DNA.**
**a**, Estimated number of monomers based on fluorescence intensity per trajectory of MeCP2 on unmethylated DNA. Bars represent mean and SD. **b**, Diffusion coefficient (*D*) for each MeCP2 trajectory on unmethylated DNA is plotted against the multimerization state of the trajectory.

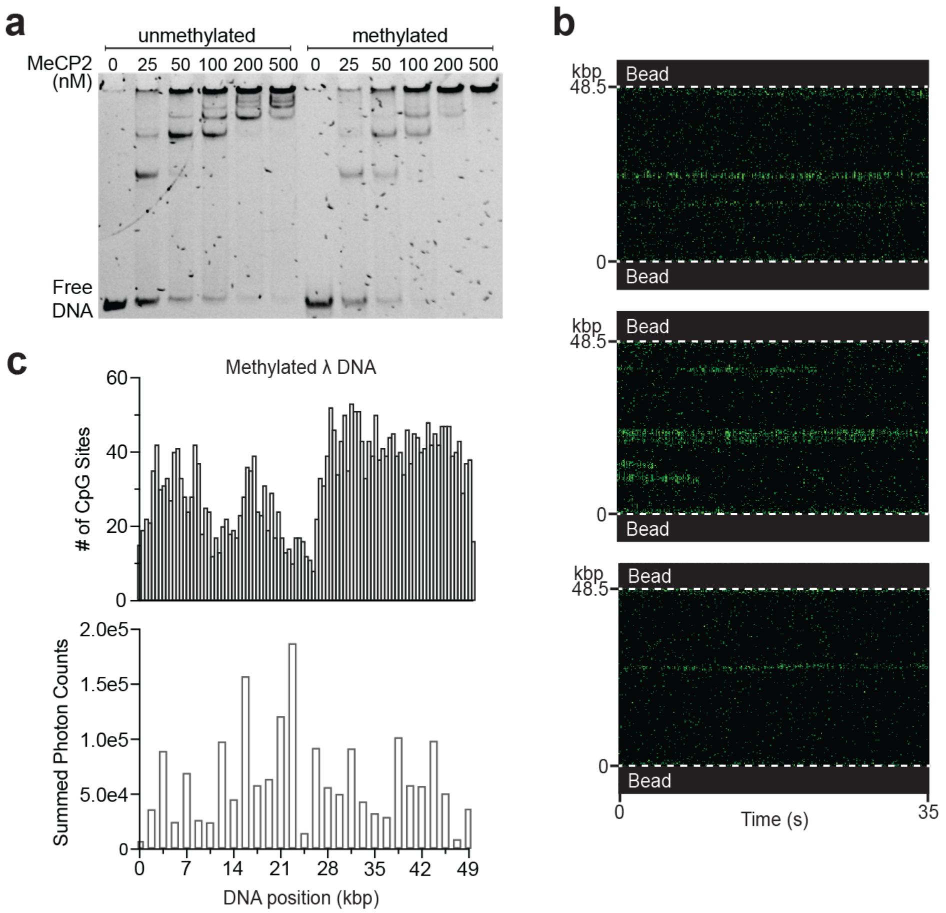


**Figure S4. Additional data of MeCP2 interaction with DNA.**

**a**, Electrophoretic mobility shift assay (EMSA) showing the products of 5 nM of 147 bp unmethylated or CpG methylated DNA incubated with an increasing concentration of MeCP2. **b**, Representative kymographs of methylated DNA tethers incubated with a low concentration (0.2 nM) of Cy3-MeCP2. **c**, (Top) Distribution of CpG sites across the length of λ DNA. (Bottom) Total Cy3-MeCP2 fluorescence signal at different DNA positions along methylated λ DNA summed over 18 tethers.

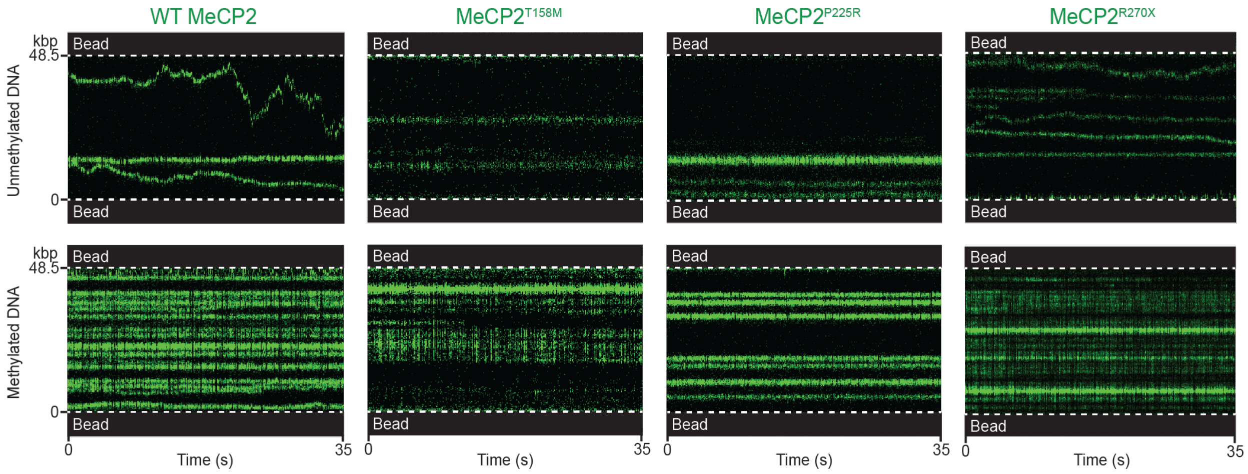


**Figure S5. Additional kymographs of DNA tethers bound with MeCP2 or RTT mutants.**

Unmethylated (top) or methylated (bottom) DNA tethers were incubated with Cy3-labeled WT MeCP2, MeCP2$^{T158M}$, MeCP2$^{P225R}$, or MeCP2$^{R270X}$.

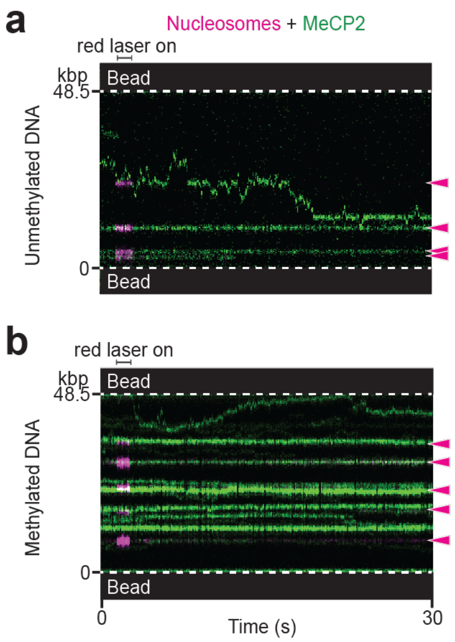


**Figure S6. Additional kymographs of nucleosome-containing DNA tethers bound with MeCP2. a**, Kymograph of an unmethylated nucleosomal DNA tether showing a rare instance where a diffusing MeCP2 trajectory crossed a nucleosome site. The other MeCP2 trajectories in this kymograph remained stationary and nucleosome-bound. Red laser was flashed on briefly to locate the nucleosomes within the tether. Arrows denote nucleosome positions. **b**, A representative kymograph of a nucleosome-containing methylated DNA tether incubated with 2 nM Cy3-MeCP2.

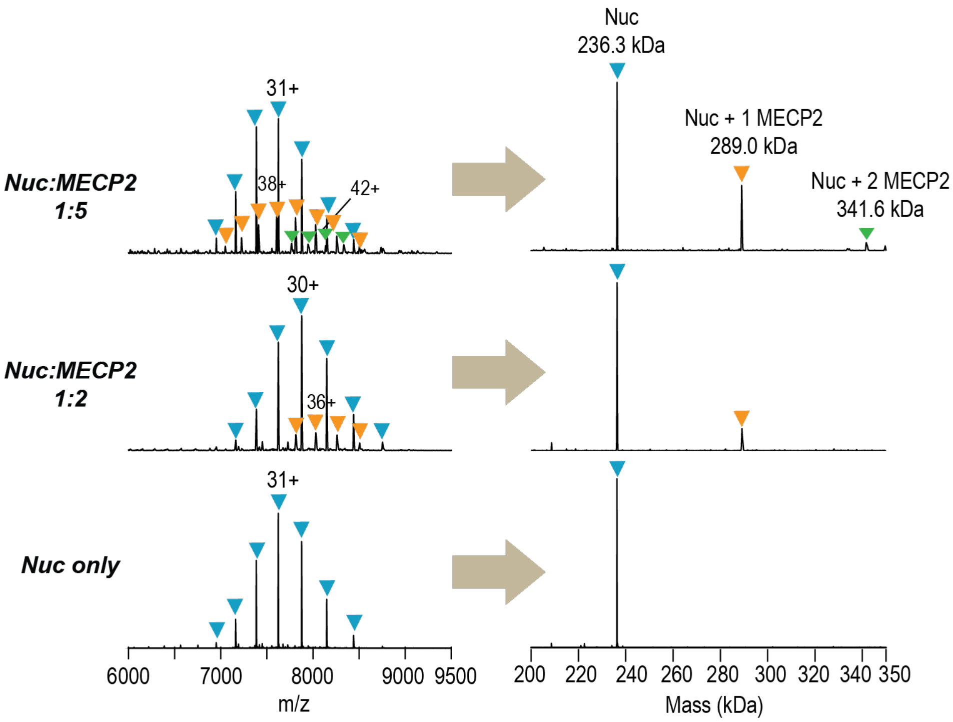


**Figure S7. Native mass spectrometry data of the MeCP2-nucleosome complex.** Native mass spectrometry spectra (left) and the corresponding deconvolved mass spectra (right) for 2 μM nucleosome incubated with different amounts of MeCP2. Addition of two-fold or five-fold molar excess of MeCP2 to the nucleosome sample yielded additional peaks corresponding to the binding of one or two 52.3-kDa MeCP2.

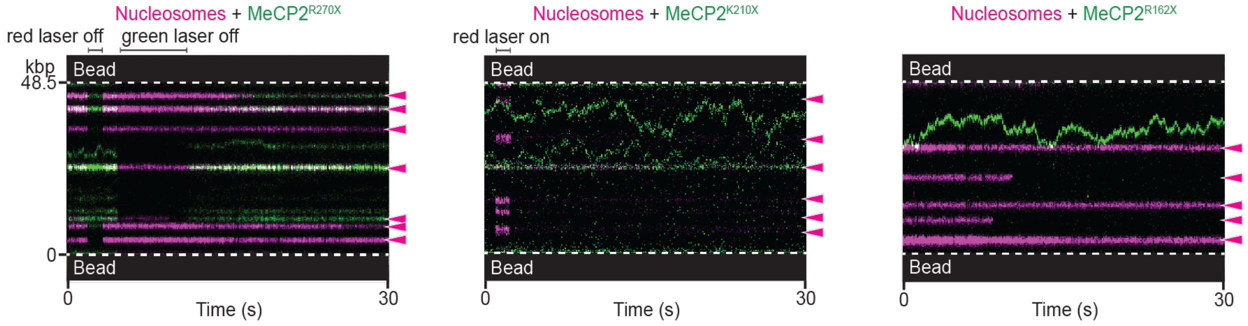


**Figure S8. Additional data of MeCP2$^{R270X}$, MeCP2$^{K210X}$, and MeCP2$^{R162X}$ binding to nucleosomal DNA.**

Representative kymographs of nucleosome-containing unmethylated DNA tethers bound with Cy3-MeCP2$^{R270X}$, Cy3-MeCP2$^{K210X}$, or Cy3-MeCP2$^{R162X}$. Arrows denote nucleosome positions.

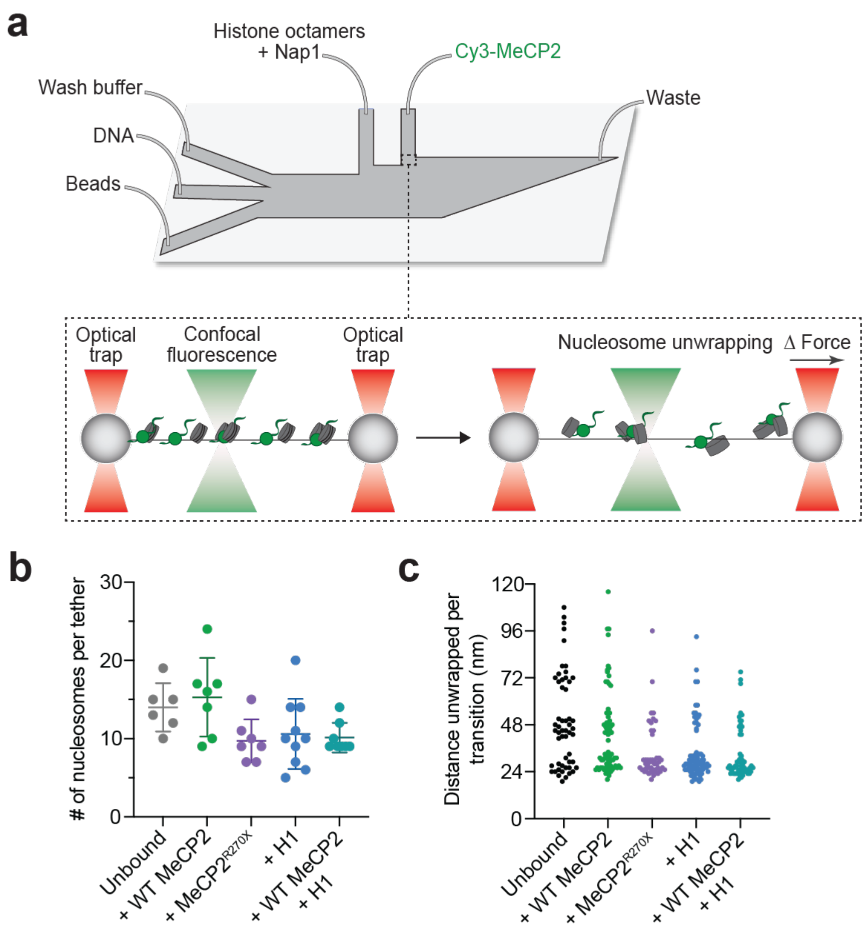


**Figure S9. Evaluation of in situ nucleosome loading on λ DNA tethers.**
**a**, Schematic of the experimental setup. Nucleosomal DNA tethers were subjected to pulling at a constant speed, resulting in force-induced nucleosome unwrapping. **b**, Number of nucleosomes loaded per tether as estimated by the number of nucleosome unwrapping events detected in the force-distance curves of tethers with no MeCP2 or H1 bound, or bound with WT MeCP2, MeCP2$^{R270X}$, H1, or with both WT MeCP2 and H1. Error bars represent SD. **c,** Distribution of the distance change per transition recorded from force-distance curves for the same conditions as in (**b**). Multiples of ~24-nm transitions confirm stochastic unwrapping of the inner turn DNA of individual nucleosomes.

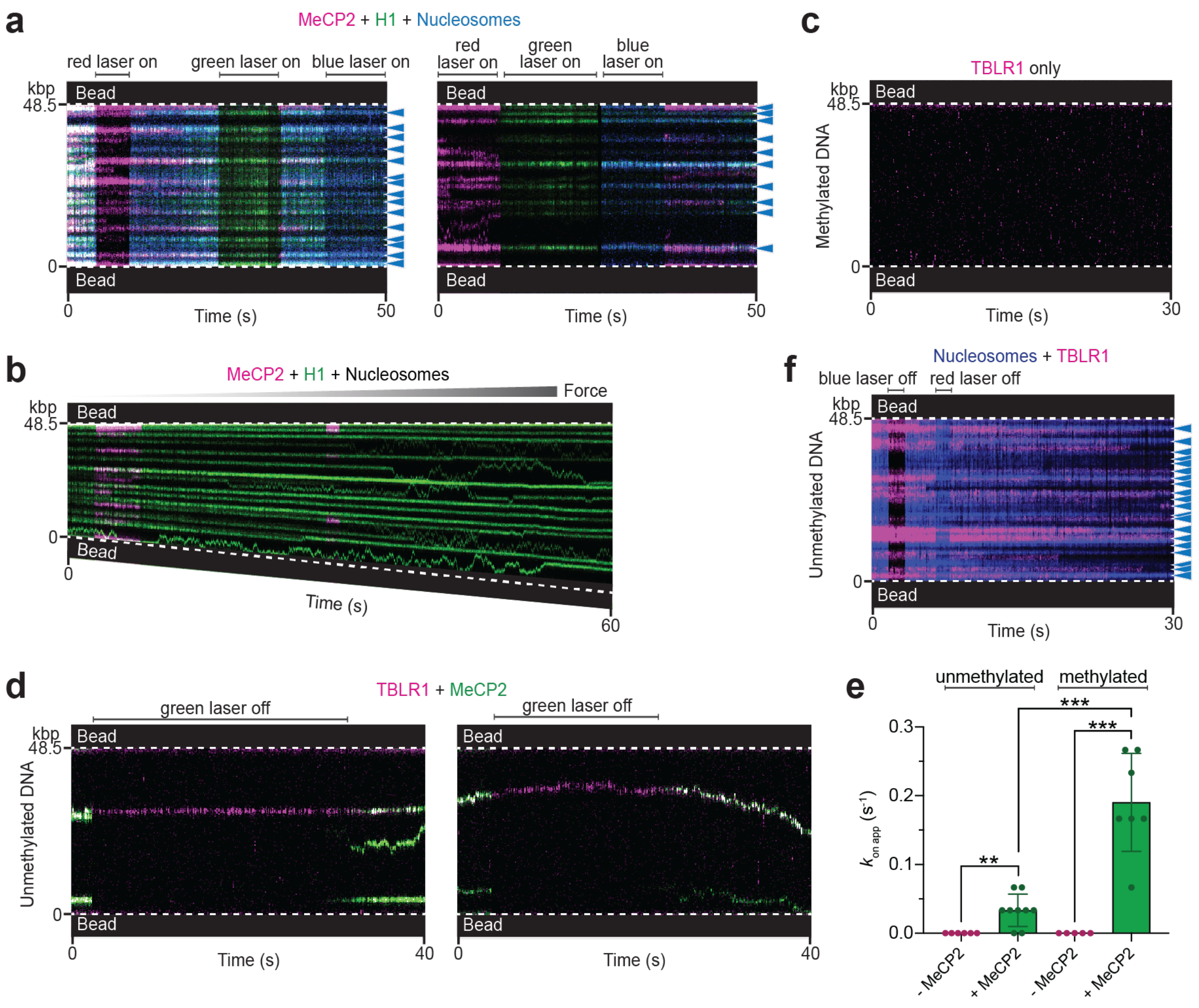


**Figure S10. Additional data of MeCP2 and its chromatin-binding partners.**
**a**, Representative kymographs of AF488-nucleosome-containing unmethylated DNA tethers bound with Cy5-MeCP2 and Cy3-H1. Arrows denote nucleosome positions. **b**, A representative kymograph of a nucleosome-containing unmethylated DNA tether bound with Cy5-MeCP2 and Cy3-H1 while being pulled to high forces. **c**, A representative kymograph of a bare methylated DNA tether incubated with LD655-TBLR1 showing lack of binding. **d**, Representative kymographs of bare unmethylated DNA tethers incubated with LD655-TBLR1 and Cy3-MeCP2. **e**, Apparent on rate for TBLR1 binding to unmethylated or methylated DNA in the absence or presence of MeCP2. Error bars represent SD. **f**, A representative kymograph of an AF488-nucleosome-containing unmethylated DNA tether incubated with LD655-TBLR1. Lasers were occasionally turned off to confirm signal from each fluorescence channel. Arrows denote nucleosome positions.
